# Method Comparison for Japanese Encephalitis Virus Detection in Samples Collected from the Indo-Pacific Region

**DOI:** 10.1101/2022.06.07.495224

**Authors:** Gary Crispell, Kelly Williams, Eric Zielinksi, Akira Iwami, Zachary Homas, Karen Thomas

## Abstract

**Introduction:** Japanese encephalitis virus (JEV) is a mosquito-borne viral pathogen, which is becoming a growing public health concern throughout the Indo-Pacific. Five genotypes of JEV have been identified. Current vaccines are based on genotype III and provide a high degree of protection for 4 of the 5 known genotypes.

**Methods:** RT-PCR, Magpix, Twist Biosciences Comprehensive Viral Research Panel (CVRP), and SISPA methods were used to detect JEV from mosquito samples collected in South Korea during 2021. These methods were compared to determine which method would be most effective for biosurveillance in the Indo-Pacific region.

**Results:** Our data showed that RT-PCR, Twist CVRP, and SISPA methods were all able to detect JEV genotype I, however, the proprietary Magpix panel was only able to detect JEV genotype III. Use of minION sequencing for pathogen detection in arthropod samples will require further method development.

**Conclusion:** Biosurveillance of vectorborne pathogens remains an area of concern throughout the Indo-Pacific. RT-PCR was the most cost effective method used in the study, but TWIST CVRP allows for the identification of over 3,100 viral genomes. Further research and comparisons will be conducted to ensure optimal methods are used for large scale biosurveillance.

## INTRODUCTION

Japanese encephalitis virus (JEV) is a mosquito-borne pathogen, which is the causative agent of Japanese encephalitis, and has become a growing public health concern throughout Asia. JEV is a flavivirus with five identified genotypes (I, II, III, VI, and V) (Li et al. 2011). JEV genotype I is currently the most prevalent genotype present throughout Asia and the Indo-Pacific. JEV genotype III was the prevailing genotype until genotype I displaced it in the 1990’s (Pan et al. 2011). The virus is thought to have originated around Indonesia and it then spread throughout Asia to include India, Japan, China, and Australia (Solomon et. al., 2003, Logibs et al., 2010, Hills et al., 2019). JEV is primarily vectored by *Culex (Cx*.*) tritaeniorhynchus*, but has been shown to be present and potentially transmitted through various other *Culex* species and *Aedes (Ae*.*) japonicus* (Oliviera et al., 2020). In fact, a thorough review of current research on JEV vector competency determined 14 species of mosquitoes to be confirmed vectors of JEV: *Ae. albopictus, Ae. vexans, Ae. vigilax, Armigeres (Ar*.*) subalbatus, Cx. annulirostris, Cx. bitaeniorhynchus, Cx. fuscocephala, Cx. gelidus, Cx. pipiens, Cx. pseudovishnui, Cx. quinquefasciatus, Cx. sitiens, Cx. tritaeniorhyncus* and *Cx. vishnui* (Auerswald et al., 2021). Vector competence relies on a multitude of factors ranging from temperature, humidity, nutrition, and also the mosquito’s midgut microbiota, immune response, and genetics (Agarwal et al., 2017). Mosquito vectors acquire the virus from animal reservoirs, mainly pigs and birds (Lindahl et al., 2012). In this fashion, the viruses found in pigs are commonly farmed, and can present risk to nearby human populations by providing the mosquito a source for acquiring the virus. Pigs show high viremia for 4-5 days after day 1 of infection with JEV and may serve as a JEV reservoir (Ricklin et al., 2016). However, there are also reports of JEV cases on Japanese islands farther away from the main island, where there are no pig farms. It is possible that wild boar are the reservoir/amplifying host (Infectious Agents Surveillance Report, 2017). In South Korea, early May to late October is the peak activity season for the *Cx. tritaeniorhynchus* species and it is when there is the greatest risk of contracting this disease (Choi et al., 2017). In a recent study, 274 *Cx. tritaeniorhynchus* mosquitos successfully fed on an animal that was infected with JEV, molecular analysis showed that 261 out of the 274 mosquitoes were found to contain JEV. The competency of *Cx. tritaeniorhynchus* has been confirmed as a JEV vector through a high infection percentage, successful virus dissemination throughout the organs, and viral particles being found in the saliva (Faizah et al., 2020).

Most reported cases of JEV infections are asymptomatic or have nonspecific febrile symptoms after 5-15 days incubation period (Amicizia et al., 2018). JEV occasionally leads to severe disease, manifesting in non-cell necrotic plaques with edema, bleeding, inflammatory infiltration in multiple regions of the brain, and severe neurologic sequelae increasing the childhood morbidity and mortality (Zhang et al., 2018). While infections rarely result in severe disease, encephalitis occurs in approximately 1 in 250 infections. Among symptomatic patients, as many as 30% will suffer fatality, and 20 to 30% of the survivors will suffer permanent disabilities such as seizures, paralysis, neurological and/or behavioral disorders. Case numbers of Japanese encephalitis (JE) vary from country to country, with the World Health Organization estimating that there are approximately 68,000 cases of JE each year world-wide. JE primarily occurs in children, with adults in endemic countries showing immunity from childhood infection. However, it is possible to contract the disease at any age. The morbidity rate is less than 1% of the infected population, but once a patient has on-set of symptoms, there is no specific treatment, and the main treatment is supportive therapy (Amicizia et al., 2018, Infectious Agents Surveillance Report, 2017). JEV prevention measures include the use of personal protective gear like insect repellents, nets, and long sleeved clothing.

Vaccination is the most effective means of preventing JEV infection (Lobigs et al., 2010). Past studies have shown that vaccination against Japanese encephalitis is an important and effective tool for prevention of disease. The cross protective capacity against other genotypes has been studied for the current Japanese encephalitis vaccine, a vaccine based on genotype III which may prevent infection for other genotypes I – IV (Elina et al., 2013). Sequence comparisons of antigenic regions of genotypes I-V indicate nucleotide (nt) similarities of less than 90% for JEV genotype I and less than 80% for JEV genotype V as compared to JEV genotype III (Sanborn et al., 2021). Serum studies have shown, JEV genotype III inactivated vaccines neutralize JEV genotype I viruses, but with reduced efficiency (Mulvey et al., 2021).

Since World War II, U.S. military personnel have been stationed in the Republic of Korea and Japan. In 1945, the islands of Heanza, Hamahika, and Okinawa had a “Japanese B type encephalitis virus” outbreak among the native population and resulted in a fatality rate of 30 percent (Sabin, 1947). It was during this time where Albert B. Sabin focused on developing a JE vaccine and successfully administered it to over 250,000 military personnel via formalin-inactivated JEV-infected mouse brains to assist in the safety of our military forces (Ratto-Kim et al., 2018). However, in 1950 there was a JEV outbreak that occurred totaling 300 cases in U.S. Soldiers stationed in Korea. This resulted in greater focus on improving the JE vaccine and developing diagnostic assays to help facilitate JE surveillance (Hoke Jr., 2005). In 2010, a JEV outbreak was recorded in the Republic of Korea (ROK) with 26 JE cases and 7 deaths reported. These numbers are likely an under-representation of the actual case count due to only severe cases being evaluated for encephalitis after hospitalization (Seo et al., 2013). Extended tours of U.S. service members and citizens in these areas greatly increased the chance of infection of JEV since U.S. personnel were typically not vaccinated against JEV (Richards et al., 2010). A 2015 study performed in the Republic of Korea tested the serum of 1,000 soldiers that had never previously deployed to determine the prevalence of JEV antibodies among personnel in the U.S. Forces Korea (Eick-Cost et al., 2015). They found that prevalence was as low as 0.2% of the population studied, and at least one soldier had titers high enough to indicate current or recent infection.

A 2018 mosquito surveillance project conducted in Camp Humphreys, ROK sequenced 6,540 Culex spp. in 260 pools. Analysis revealed 122 distinct virus species in the pools (Sanborn et. al., 2021). Two of these pools were positive for JEV genotype V along with other viruses. Discovery of JEV genotype V in a highly populated area adjacent to Seoul is of concern, due to questions regarding vaccine effectiveness against JEV genotype V. Currently, it is recommended by the U.S. Centers for Disease Control and Prevention that travelers to JE-endemic countries receive a 2-dose series of IXIARO prior to travel, particularly for those travelers with longer plans (greater than a one month stay), those who will frequently travel to JEV-endemic countries, or those with shorter stays that may be at greater risk due to planned activities, season of travel, or type of accommodation during stay (CDC, 2019). Given the potential serious outcomes and effect on force readiness, it is of great concern for the public health of both military and civilian populations to accurately detect JEV to ensure the safety of our military populations.

Viral encephalopathies continue to be a public health concern in many parts of the world. In this study, we examine and compare various detection methods for Japanese encephalitis virus to determine which are the most sensitive, cost effective, and provide the most data to the investigating scientists. The methods covered are RT-PCR, Luminex MagPix technology, the TWIST Comprehensive Viral Research Panel, direct MinION sequencing, and a SISPA method utilized with both the MinION and MiSeq.

## MATERIALS AND METHODS

### Mosquito Identification and Grouping

Mosquito samples were identified and sorted by an entomologist according to collection date, collection site, and species. Specimen pools consisted of varying numbers of mosquitoes (1 to 30) in each microcentrifuge tube. Collections that exceeded more than 30 specimens were sorted in to multiple tubes. Mosquito pools were maintained at -80°C until the time of homogenization and nucleic acid isolation.

### Materials

All common laboratory supplies and reagents were obtained through Thermo Fisher Scientific (Waltham, MA, USA) and its subsidiaries unless specifically noted. Molecular grade isopropanol and ethanol were sourced locally through Kanto Chemical Co. (Chuo-ku, Tokyo, Japan).

### Nucleic Acid Isolation

Frozen mosquito samples stored in microcentrifuge tubes were allowed to thaw at room temperature for up to 15 minutes. Using the Zymo Direct-zol™-96 MagBead RNA kit (Zymo Research, Irvine, CA, USA), 400 μL of TRI reagent and two RNase-free 3.2mm stainless steel beads (Next Advance, Troy, NY, USA) were added to each specimen tube. Samples were lysed using the Qiagen TissueLyser II for 7 minutes at a frequency of 24/s followed by centrifugation for 10 minutes at 14,000 rcf. 200 μl of supernatant was removed from each sample tube and pipetted into a 96-deep well plate. 200μL of 99.5% ethanol, 20μL of MagBinding beads, and 5μL of Proteinase K were added to each sample well before adding the plate to the Thermo Scientific KingFisher Flex automated instrument. An extraction control, consisting of all reagents and no mosquito matrix, was added to the sample plate as well. The elution plate was stored at -20°C for short-term storage or used immediately for further analysis on the Applied Biosystems ABI 7500 Fast DX, MagPix, MinION and MiSeq. A second extraction was performed on the homogenized mosquitos with the addition of 50μL DNase added to the sample prior to being placed on the Thermo Scientific KingFisher Flex automated instrument.

### Nucleic Acid Quantification

Isolated nucleic acid was quantified on the Qubit 3.0 Fluorometer to evaluate isolation procedures. Following the Thermo Fisher Scientific Qubit 3.0 Fluorometer BR (broad range) RNA protocol, a low and high nucleic acid standard were prepared and analyzed first to ensure quality before samples were tested.

### Semi-Quantitative RT-PCR

Semi-Quantitative RT-PCR was performed within the specifications provided by the Applied Biosystems TaqMan Fast Virus Master Mix protocols. Frozen reagents and nucleic acids were thawed on ice. The mastermix formula used was 5μL of TaqMan Fast Virus, 1 μL of the 10μM gene-specific forward and reverse primers, 0.80 μL of gene-specific probe at 5μM (Table 1) and 7.20μL of nuclease free water per reaction. Each reaction consisted of 15μL of mastermix along with 5μL of nucleic acid. An extraction control, non-template control, and positive control were used for each plate. The following thermocycling conditions were used on the ABI 7500 Fast Dx instrument: 50°C for 5 min; 95°C for 20 s; 40 cycles of 95°C for 3 s and of 60°C for 30 s. The amplification plot and Ct values were used to determine the presence of Japanese Encephalitis Virus (JEV). Viral copies were estimated using a standard curve formed from a synthetic JEV control ordered from Integrated DNA Technologies (IDT). It was calculated that 2.4 quadrillion copies of the control were present in 5μL of the undiluted control based on data provided on the material sheet submitted with the control from IDT. Serial dilutions were run in duplicate, and the average of the two runs were plotted against the log of the estimated copies used in each run. A standard curve was used to determine the number of copies in each of the serial dilutions of the JEV synthetic plasmid control with an R^2^ value of 0.9895. Semi-quantitation was carried out by extrapolating used Ct values of positive samples against the standard curve (**Supplementary Figure 1**).

**Table 1.**
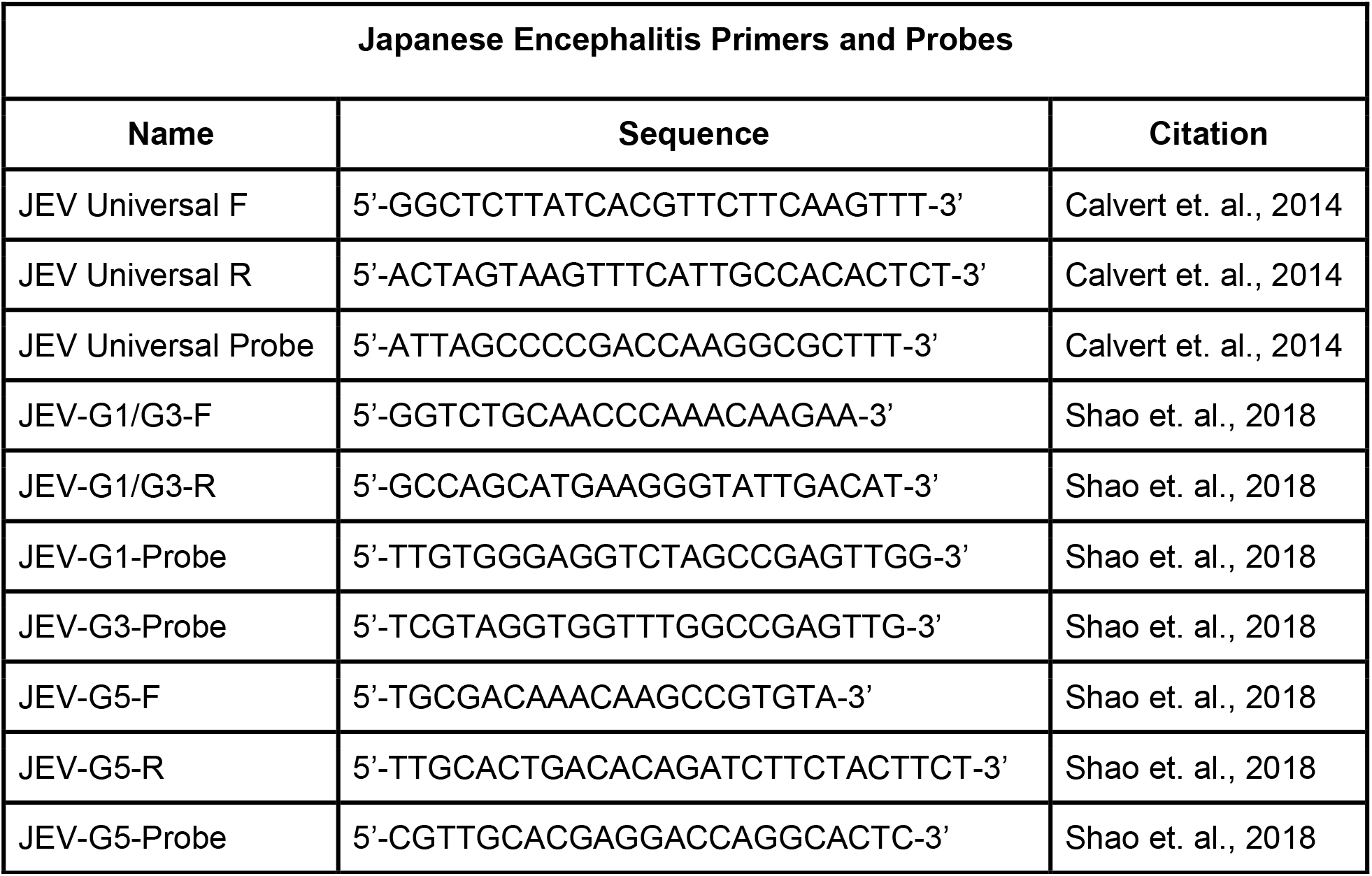
Primers and probes used to detect and genotype JEV in this study.

### MagPix Mega Mosquito Panel

Nucleic acids from positive samples, as detected by RT-PCR, were reacted in accordance to the instructions for use included with the GenArraytion Inc. MultiFLEXMega Mosquito Borne Panel (South Orange, NJ, USA). In the first step, RNA was converted into cDNA before PCR utilizing the Qiagen One-Step PCR Kit and panel-specific primers according to the instructions included with the panel. Additionally, five different controls from GenArraytion were run with the samples to ensure that all targets amplified as expected. Next, 10μL of the PCR products was mixed with 5μL of panel-specific beads and 35μL Buffer A and beads were hybridized to the target DNA/cDNA according to instructions. Finally, the beads were placed on a magnetic plate and liquid removed via pipette. The beads were quickly taken off the magnetic plate and a streptavidin solution of 5μL SAPE and 70μL Buffer B was mixed with the beads and heated at 52°C to allow the beads to fluoresce in the MagPix instrument. Plates were maintained at 52°C and quickly entered onto the pre-heated instrument so that analysis could be conducted.

### TWIST CVRP

Total nucleic acid library preparation and target enrichment standard hybridization workflow followed the guidelines of Twist Next Generation Sequencing (NGS) protocols. Three previously extracted nucleic acid samples were diluted along with synthetic controls. cDNA synthesis and purification followed. Next, DNA fragmentation, telomere repair, and dA-Tailing were performed. Universal Twist adapters were then ligated to the cDNA and purified. Finally, PCR amplification was conducted to index the samples and finish the library preparation portion of the study. A single pooled library was first prepared from the indexed library-prepped sample pools. This was followed by hybridization of the targets in solution, which was approximately 16 hours in total to complete. Next, the binding of hybridized targets to desired streptavidin beads occurred. Libraries were then enriched via PCR amplification and purification. Sample libraries were ready for sequencing on the Illumina NGS platform according to manufacturer protocols after PCR amplification and purification. Sequencing data was processed on the One Codex bioinformatics online platform and according to methods described below.

### Sequence-Independent, Single-Primer-Amplification Mediated MiSeq Sequencing

Sequence-Independent, Single-Primer-Amplification (SISPA) is a method of tagged random amplification of nucleic acid targets that has been shown to be suitable for preparing samples for whole genome viral sequencing (Wright et. al., 2015). Random hexamer tagging with 20 nt barcode “K” (GACCATCTAGCGACCTCCAC) was performed using primer K8N (GACCATCTAGCGACCTCCACNNNNNNNN) as described by Chrzastek et al in 2017. Library preparation was performed with a protocol provided by USAMRIID Center for Genome Sciences using the K/K8N primers as described above. In short, first strand synthesis is performed with primer K8N, dNTPs, RNA template and nuclease-free water. This is heated at 65°C for 5 minutes, placed on ice and then 5X SuperScript IV buffer, DTT, RNaseOUT Recombinant RNase Inhibitor are added to each library before incubating for 10 minutes each at 23°C, 50°C and 80°C and held at 4°C. Second strand synthesis was performed with RNAseH and Klenow 3’ -> 5’ Exo DNA polymerase added to the first strand mix and incubated for 30 minutes at 37°C, 20 minutes at 75°C and held at 4°C. Cleanup was performed with AMPureXP beads according to protocol. Random fragment amplification was performed with primer K, 5x Phusion HF buffer, dNTPs, nuclease-free water, and Phusion Hot Start II DNA polymerase and template cDNA. This mix was cycled for 30s at 98°C, 40 cycles of 10s at 98°C, 10s at 50°C, and 45s at 72°C, and a final cycle of 72°C for 10 minutes before being held at 12°C. A new mix was made and added to each sample for a final cycle of 30s at 98°C, 10s at 50°C, and 72°C for 10 minutes and held at 12°C. PCR products were then cleaned using the AgenCourt Ampure XP Beads before moving to finishing library prep and loading on the MiSeq. PCR products were fragmented, tagged with adaptors, and appended with unique dual index sequences via secondary PCR using reagents from the NEBNext Ultra II Directional RNA Library Prep Kit for Illumina and NEBNext Multiplex Oligos for Illumina (Dual Index Primers Set 1) according to manufacturer protocol.

### MinION sequencing

Two methods of minION sequencing were attempted. Unprocessed whole nucleic acid extract as well as SISPA mediated processing of nucleic acid as described above were used to generate cDNA. The cDNA was then processed using the Nanopore Rapid Sequencing Kit SQK-RAD004 for library preparation. 7.5μl of undiluted cDNA was used in the library preparation process to obtain as near to 400ng of cDNA as possible. The rest of library prep and loading onto the minION was done according to manufacturer procedure.

### Bioinformatics

Sequence data was treated based on single or paired reads and aligned to target genomes and visually mapped with Tablet software (The James Hutton Institute, Scotland, UK). MinION FASTQ single reads were combined for single file processing using the concatenate (cat) command and FASTA files were generated using seqtk (Heng, 2018). MiSeq paired read data were first joined using FLASH (Fast length adjustment of short reads) for single file processing (Magoc, T, Salzberg, S. L., 2011). FASTQ files were then fed into the NanoPipe and assembled and aligned against target genomes (Shabardina et. al., 2019). Tablet software was used to view the resulting BAM files mapped against the target genome for visual coverage of the genome and mismatch data, see **Supplementary Figures 2 and 3**. (Milne et. al., 2013).

## RESULTS

Of 549 mosquito pools, 21 pools were found to be positive, corresponding to a 3.8% positivity rate from *Cx. tritaeniorhynchus* as detected using described methods. A majority of the positive samples analyzed using the ABI 7500 Fast Dx platform and Fast Virus RT-PCR mastermix provided us with relatively low Ct values upon extraction (**Table 2)**. Analyzing the positive samples with Qubit provided total RNA concentrations per sample. The Broad Range Qubit Assay was suitable for running samples with Ct values up to 20.0, and High Sensitivity Assay should be considered for samples with higher Ct values. Utilizing the primers and probes as shown in **Table 1**, the samples were then genotyped via RT-PCR. **Table 2** shows that all 21 JEV positive samples produced sigmoidal curves when tested for JEV genotype I and none of the samples produced any curve when tested for JEV genotype III or V. The positive control used did not produce a sigmoidal curve for JEV genotype I, but did so for genotype III.

**Table 2.** JEV positive samples with their respective Ct values, and genotype. G1-genotype I, G3 - genotype III, G5 - genotype V.

| Sample | Species | # in Pool | Ct | JEV G1 | JEV G3 | JEV G5 |
| --- | --- | --- | --- | --- | --- | --- |
| A21.2333 | *Cx. tritaeniorhynchus* | 21 | 16.79 | ✓ | X | X |
| A21.2437 | *Cx. tritaeniorhynchus* | 30 | 18.61 | ✓ | X | X |
| A21.2560 | *Cx. tritaeniorhynchus* | 30 | 18.61 | ✓ | X | X |
| A21.2711 | *Cx. tritaeniorhynchus* | 30 | 34.99 | ✓ | X | X |
| A21.2838 | *Cx. tritaeniorhynchus* | 30 | 20.29 | ✓ | X | X |
| A21.2886 | *Cx. tritaeniorhynchus* | 30 | 18.67 | ✓ | X | X |
| A21.3160 | *Cx. tritaeniorhynchus* | 30 | 24.52 | ✓ | X | X |
| A21.3171 | *Cx. tritaeniorhynchus* | 30 | 20.53 | ✓ | X | X |
| A21.3232 | *Cx. tritaeniorhynchus* | 19 | 18.82 | ✓ | X | X |
| A21.3389 | *Cx. tritaeniorhynchus* | 30 | 33.84 | ✓ | X | X |
| A21.3401 | *Cx. tritaeniorhynchus* | 30 | 24.29 | ✓ | X | X |
| A21.3402 | *Cx. tritaeniorhynchus* | 30 | 19.48 | ✓ | X | X |
| A21.3404 | *Cx. tritaeniorhynchus* | 30 | 30.55 | ✓ | X | X |
| A21.3465 | *Cx. tritaeniorhynchus* | 30 | 19.10 | ✓ | X | X |
| A21.3577 | *Cx. tritaeniorhynchus* | 30 | 20.08 | ✓ | X | X |
| A21.3585 | *Cx. tritaeniorhynchus* | 30 | 29.63 | ✓ | X | X |
| A21.3587 | *Cx. tritaeniorhynchus* | 30 | 18.12 | ✓ | X | X |
| A21.3675 | *Cx. tritaeniorhynchus* | 30 | 17.70 | ✓ | X | X |
| A21.3678 | *Cx. tritaeniorhynchus* | 30 | 23.29 | ✓ | X | X |
| A21.3682 | *Cx. tritaeniorhynchus* | 30 | 23.31 | ✓ | X | X |
| A21.3683 | *Cx. tritaeniorhynchus* | 30 | 18.62 | ✓ | X | X |
| JEV 1:1,000 | Unknown | N/A |  | X | ✓ | X |

In total, 21 JEV positive sample pools and 21 JEV negative sample pools, were analyzed with the MagPix instrument using the GenArraytion Mega Mosquito-borne MultiFLEX® Panel to assess for the presence of JEV, in addition a historical sample used as a RT-PCR control at a dilution of 1:1,000 was analyzed. The data in **Figure 1 and Supplementary Table 1** show that the Net MFI was below 200, a negative result, for all JEV samples tested, with the positive control MBP_CD over 4,000 and the historic RT-PCR control for JEV at a 1:1,000 dilution was positive at over 500. A signal above 300 is considered a positive result for each analyte and below 200 is negative, per manufacturer’s insert. All Internal, Fluorescence and Non-Specific Binding Controls were acceptable for samples and controls ran.

**Figure 1.**
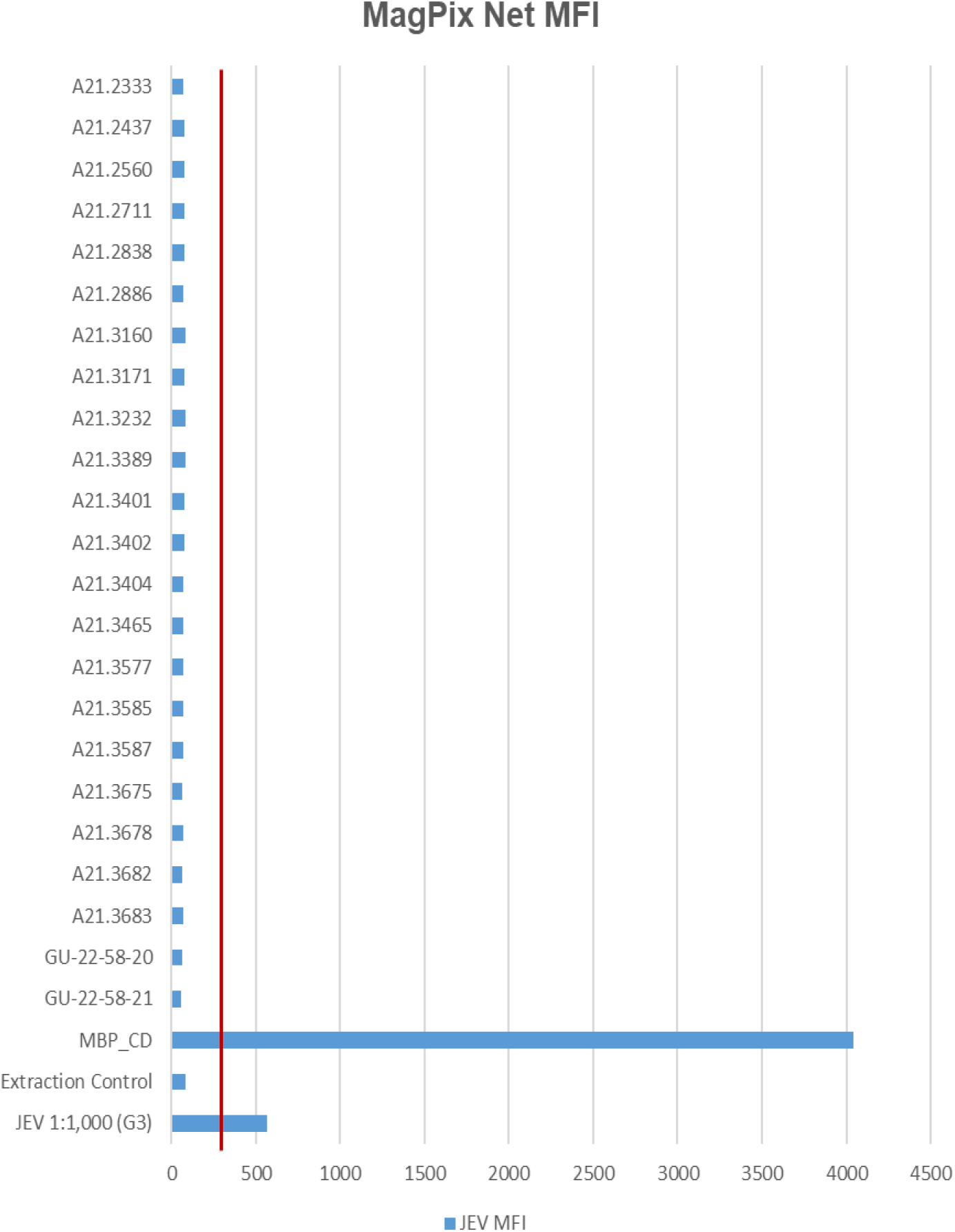
MFI values for MagPix MultiFLEX® Panel analytes. Red line represents 300 MFI; samples above 300 MFI are positive. Samples labeled A21.xxxx are positive for JEV as determined by RT-PCR. Samples labeled GU-22-58-xx are negative for JEV. JEV 1:1,000 (G3) is an archived sample. Internal Control, Fluorescence Control and Non-specific Binding Control had comparable values between specimen types. (MFI = Median Fluorescent Intensity; MBP_CD

Three (3) JEV positive samples were sequenced using the Twist Comprehensive Viral Research Panel (CVRP) on the Illumina MiSeq. All three samples were confirmed to contain JEV via sequencing, and additionally a mosquito specific virus, the Yichang virus, was detected in two of the three samples. One sample, sample 3171, showed evidence of fungal contamination with the One Codex platform detecting the presence of Cryptococcus neoformans. MiSeq FASTQ files were also assembled and aligned to JEV Genotypes I (GenBank Accession JN381833.1), III (GenBank Accession KP164498.2), and V (GenBank Accessions HM596272.1 and JF915894.1). Comparisons indicated significant alignment to Genotype I versus III and V with a Kruskal-Wallis test of mismatch percentage of each type showed a p-value of 0.009343. This information and the relative abundance are summarized in **Table 4** and **Figure 2** below.

**Table 4.** Bioinformatics Results of TWIST CVRP and SISPA Sequencing Data. Sample names and mismatch percentage for each genotype. *MiSeq Pool 1 results are estimates due to low number of aligned reads.

| Sample | JEV G1 | JEV G3 | JEV G5 Muar | JEV G5 XZ0934 |
| --- | --- | --- | --- | --- |
| **Twist Mismatch Percentages** |  |  |  |  |
| **A21.2333** | 2.7 | 9.3 | 17.1 | 17.4 |
| **A21.2886** | 3.1 | 10.7 | 18.9 | 19.0 |
| **A21.3171** | 3.4 | 10.2 | 18.0 | 18.2 |
| **SISPA Mismatch Percentage** |  |  |  |  |
| **MinION Pool 1** | 14.4 | 19.9 | 21.7 | 23.1 |
| **MinION Pool 2** | 12.6 | 19.4 | 22.4 | 22.6 |
| **MiSeq Pool 1** | 2.7* | 9.3* | 18.3* | 17.7* |

**Figure 2.**
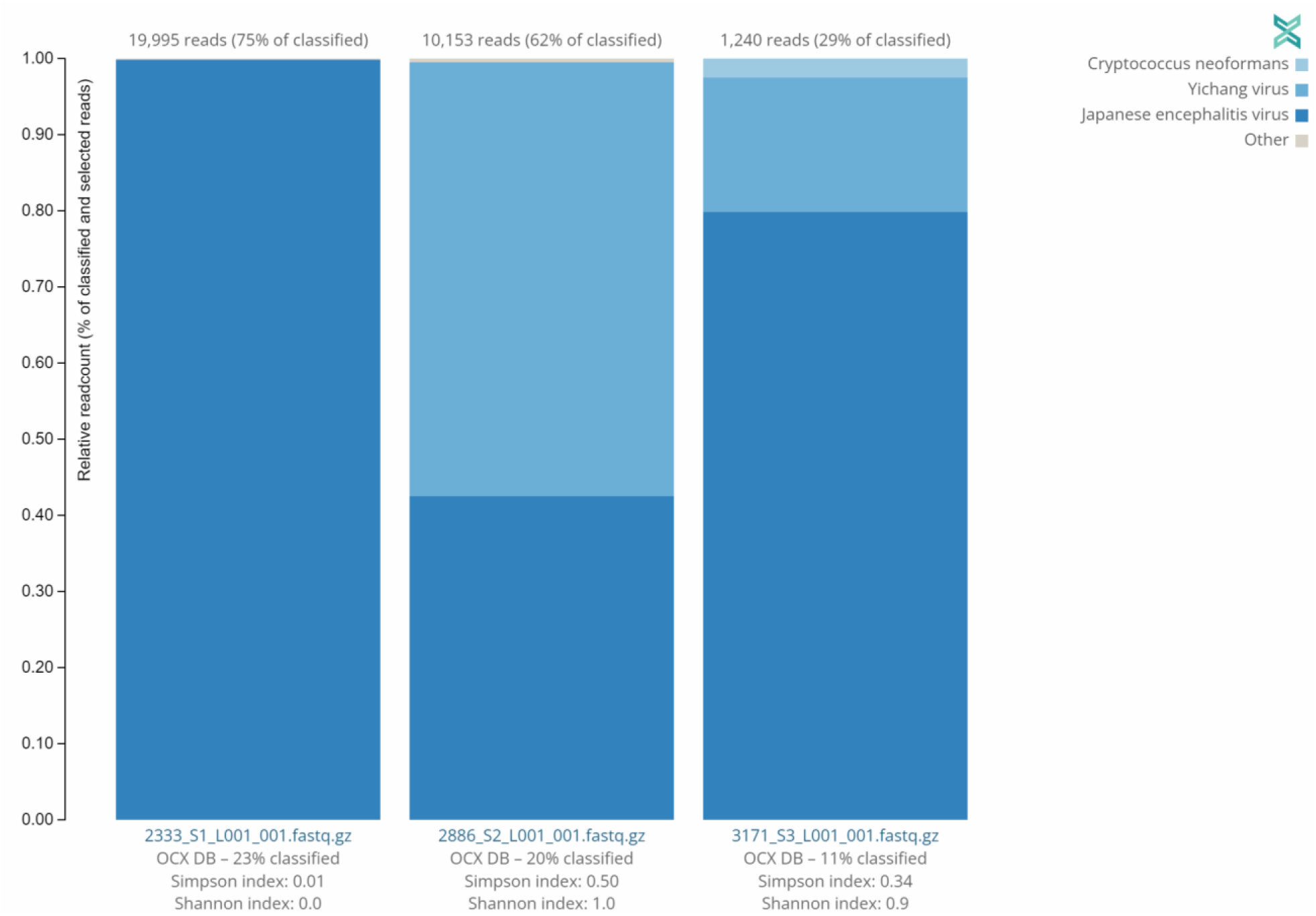
Distribution of sequences identified by Twist CVRP. Three of the JEV positive samples analyzed with the Twist Biosciences CVRP are shown with their read counts and individual distributions of Japanese encephalitis virus, Yichang virus, *Cryptococcus neoformans*, and other sequences.

MinION sequencing of unprocessed total nucleic acid provided ample coverage of the Cx. tritaeniorhynchus mitochondrial genome, however, it was not able to provide any reads necessary to determine that JEV was present in the sample when using unprocessed nucleic acid for direct sequencing. SISPA mediated sequencing on the minION was able to provide near full genome coverage using nanopipe. Analysis of the various genotypes (G1, G3, G5) showed agreement with Twist results, but with a higher mismatch percentage (**Table 4**). The resulting reads from the SISPA sequencing on the MiSeq were combined and processed. Very few reads were aligned to the JEV genomes, so accuracy of the mismatches can’t be assessed, however they appear to match Twist results (Table 4).

## DISCUSSION

RT-PCR was the quickest, easiest, and most cost-effective method for detection of JEV in our study, requiring less than 2 hours from setup to results. The ABI 7500 provided accurate and reproducible results for both genotypes I and III, while the MagPix Mega Mosquito panel was able to detect JEV genotype III from an archived sample, but not JEV genotype I in our recently extracted mosquito samples. Due to lack of Genotype II, IV, and V sample material, it has yet to be determined if the panel can detect the remaining JEV genotypes. For proper public health surveillance, our lab requires the ability to test for and detect all genotypes of JEV. Currently, the Mega Mosquito panel does not instill confidence to support the Public Health Command - Pacific’s goal of vector-borne disease detection in high risk areas where JEV infection is possible, especially when considering lower throughput, increased analytical time compared to RT-PCR, and ten-times the analysis cost per pool. Because of these factors combined, future surveillance will be conducted using the RT-PCR centric approach and genotype identifications will be carried out using NGS based solutions.

This study focused primarily on the detection of JEV types I, III, and V as the majority of our arthropod samples originated in Eastern Asian countries and the Pacific Islands. There is conflicting data regarding if the current Japanese encephalitis vaccine, derived from G3 JEV, can induce protective immunity against the remaining genotypes. Furthermore, vaccine potency against the emerging G5 genotype has not yet been reported (Cao et. al., 2016). More detailed studies demonstrating increased levels of protective antibodies against JEV Genotype V are needed. Research suggests that JEV genotype V re-emerged after nearly a half century hiatus. The first strain or the Muar strain was isolated in Malaya in 1952. The second was extracted from mosquito samples collected in China in 2009 and designated strain XZ0934. The emergence of JEV genotype V has been detected in the Republic of Korea (ROK) as recently as 2016-2018. A JEV genotype shift may be occurring in the ROK. Initially, the prevalent JEV genotype was identified as Genotype III until approximately 1990s when it shifted to Genotype I, and it may be shifting once again to Genotype V, which was first identified in 2010 (Sanborn et al., 2021). Therefore, delivering optimal surveillance detection methods against all predominant JEV genotypes in the surrounding area is critical for the overall well-being of both civilian and military personnel.

Identifying predominant genotypes by area will allow public health officials to make informed and more effective decisions concerning prevention, testing, and vaccine allocation/development. Utilizing the real-time PCR with universal and genotype specific primers/probes (**Table 1)**, we successfully genotyped all positive samples as JEV genotype I as shown in **Table 2**. This supports current data indicating JEV genotype I is the predominant genotype within our testing areas of the Indo-Pacific. We further confirmed findings using the Twist Comprehensive Viral Research Panel on the Illumina MiSeq platform. OneCodex software provided for use with the Twist panel detected JEV, but did not provide typing information. However, we were able to use the sequencing data generated from the protocol to analyze via our own pipeline. Using a Kruskal-Wallis Rank Sum Test comparing the amount of aligned base pairs from each sample to each genotype, provided alignment data that suggested JEV genotype I. With a p-value of 0.009343, the differences in alignment between Type I and Types III and V were highly significant. This method is certainly useful for pathogen discovery, however, given the multiple days and hours of hands on bench time required as well as the very significant cost of reagents required for the protocol, we cannot recommend this method for routine screening for pathogens. SISPA library preparation has the benefit of requiring less bench time to sequencing compared to the Twist CVRP protocol. Additionally, we found that the MinION’s capability for long sequencing reads, was able to provide better coverage of the genome overall as compared to our attempt at SISPA on the MiSeq platform. Given the relative ease of library preparation, SISPA sequencing on the MinION may be a less expensive alternative that is capable of field deployment. The downside is that there can be a higher rate of errors in reads when using the minION platform. While more data is required to determine the sensitivity of our genotyping RT-PCR assay, it seems likely that RT-PCR will be sufficient for providing not only screening of JEV but also typing of positive samples.

## CONCLUSION

This study demonstrates that utilization of the Direct-zol-96 MagBead RNA kit with the KingFisher Flex for automated nucleic acid isolation coupled with RT-qPCR techniques on the ABI 7500 remain the most sensitive, fastest, and cost-effective method for detection and genotyping of JEV in *Cx. tritaeniorhynchus* samples. These methods are especially useful when processing large batches of samples. For broader passive pathogen discovery, other sequencing based techniques such as SISPA or Twist panels may be appropriate depending on the goal of the laboratory, as these methods do not require target specific reagents and interpretation of results is only limited by the availability of viral genomic data. SISPA in particular has the potential to be performed in the field for passive detection of novel or emerging pathogens, but suffers from the lack of sequencing depth acquired through random amplification. The Twist panel shows greater promise for broad detection and potential genotyping despite the extended bench time required to prepare and sequence the hybridized libraries.

## DISCLAIMER

The views expressed in this presentation are those of the authors and do not necessarily reflect the official policy of the Department of the Army, Department of Defense, or the U.S. Government.

## ACKNOWLEDGEMENTS

We would like to thank Dr. Craig Stoops of the 65^th^ Medical Brigade at USAG Camp Humphreys, South Korea for his efforts in trapping and identifying the mosquitoes used in this study. We would also like to thank entomologists CPT John J. Eads and Mr. Kei Jimbo for receiving, processing, and submitting the mosquitoes used in these findings. Special thanks to USAMRIID’s Jeffery Kugelman and Nicholas Di Paola for providing the SISPA protocol and for the hands-on laboratory training.

## SUPPLEMENTARY MATERIALS

**Supplementary Figure 1.**
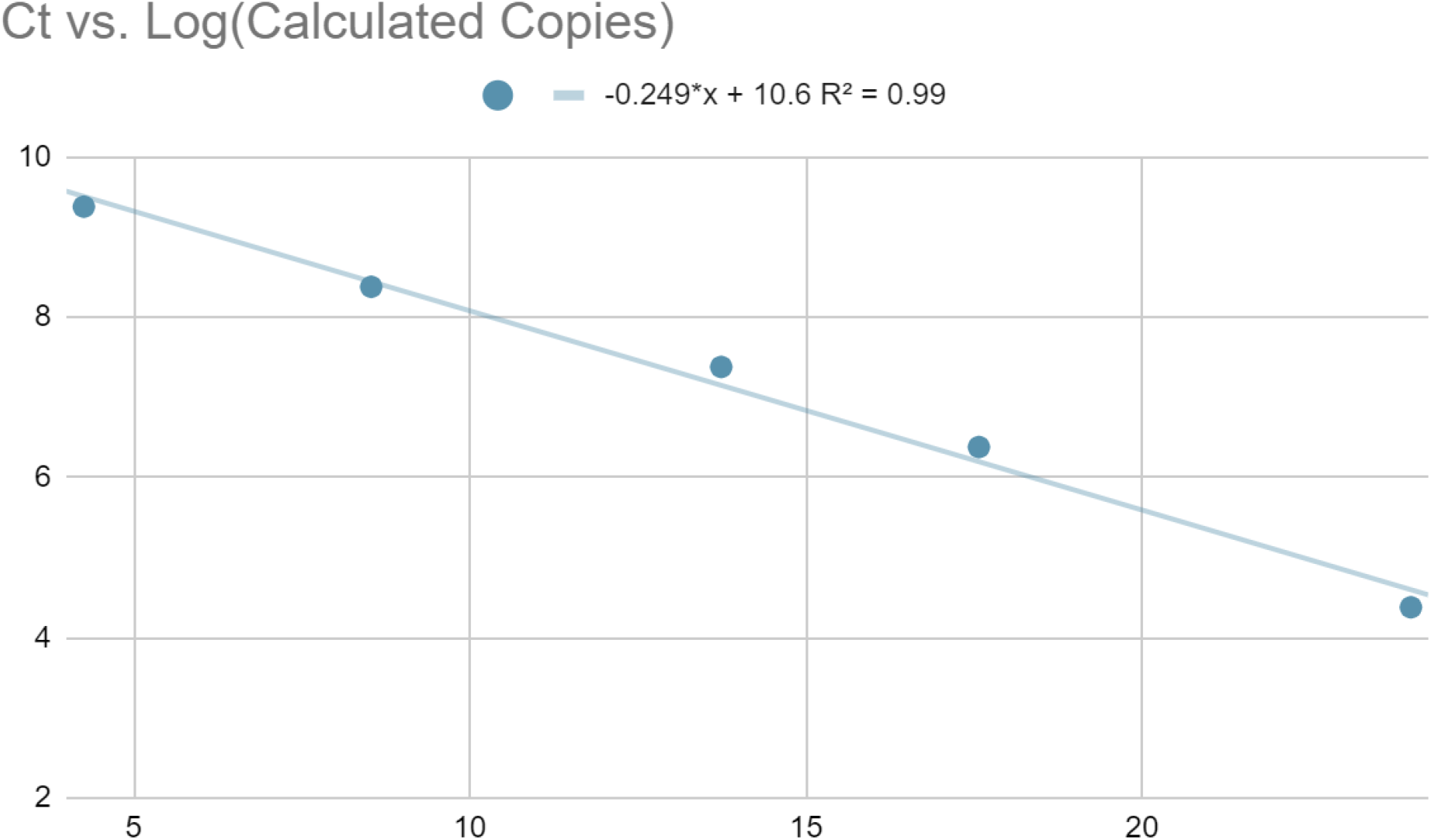
Ct values of serial dilutions of synthetic JEV control. Ct values were plotted against the log value of the calculated copy number of control used in each run.

**Supplementary Table 1.** MagPix Mega Mosquito-borne Pathogen MultiFLEX® Panel analytes and their respective net MFI values. (JEV - Japanese Encephalitis; IC - internal control; FC - fluorescence control; NSBC - non-specific binding control)

| MagPix Net MFI |  |  |  |  |
| --- | --- | --- | --- | --- |
| Sample | JEV | IC | FC | NSBC |
| A21.2333 | 68 | 5576 | 3259 | 68 |
| A21.2437 | 79 | 5543 | 3065 | 75 |
| A21.2560 | 75 | 5606 | 3292 | 76 |
| A21.2711 | 75 | 5527 | 3615 | 75 |
| A21.2838 | 77 | 5575 | 3472 | 79 |
| A21.2886 | 72 | 5550 | 3239 | 74 |
| A21.3160 | 80 | 5560 | 3917 | 79 |
| A21.3171 | 78 | 5522 | 4333 | 74 |
| A21.3232 | 84 | 5570 | 5331 | 78 |
| A21.3389 | 82 | 5531 | 5109 | 86 |
| A21.3401 | 75 | 5555 | 3820 | 77 |
| A21.3402 | 75 | 5490 | 3612 | 76 |
| A21.3404 | 73 | 5560 | 3474 | 76 |
| A21.3465 | 69 | 5580 | 3362 | 75 |
| A21.3577 | 71 | 5515 | 3010 | 77 |
| A21.3585 | 72 | 5597 | 3121 | 73 |
| A21.3587 | 72 | 5516 | 4204 | 74 |
| A21.3675 | 65 | 5556 | 3213 | 71 |
| A21.3678 | 71 | 5526 | 3027 | 74 |
| A21.3682 | 64 | 5484 | 2760 | 74 |
| A21.3683 | 68 | 5500 | 2651 | 71 |
| GU-22-58-20 | 62 | 5498 | 3631 | 62 |
| GU-22-58-21 | 59 | 5567 | 2746 | 60 |
| MBP_CD | 4041 | 5607 | 3059 | 91 |
| Extraction Control | 81 | 5567 | 4127 | 85 |
| JEV G3 1:1,000 | 564 | 5654 | 2892 | 80 |
| MBP_CD | 8332 | 5655 | 8795 | 98 |
| Extraction Control | 97 | 5542 | 9619 | 84 |

**Supplementary Figure 2.**
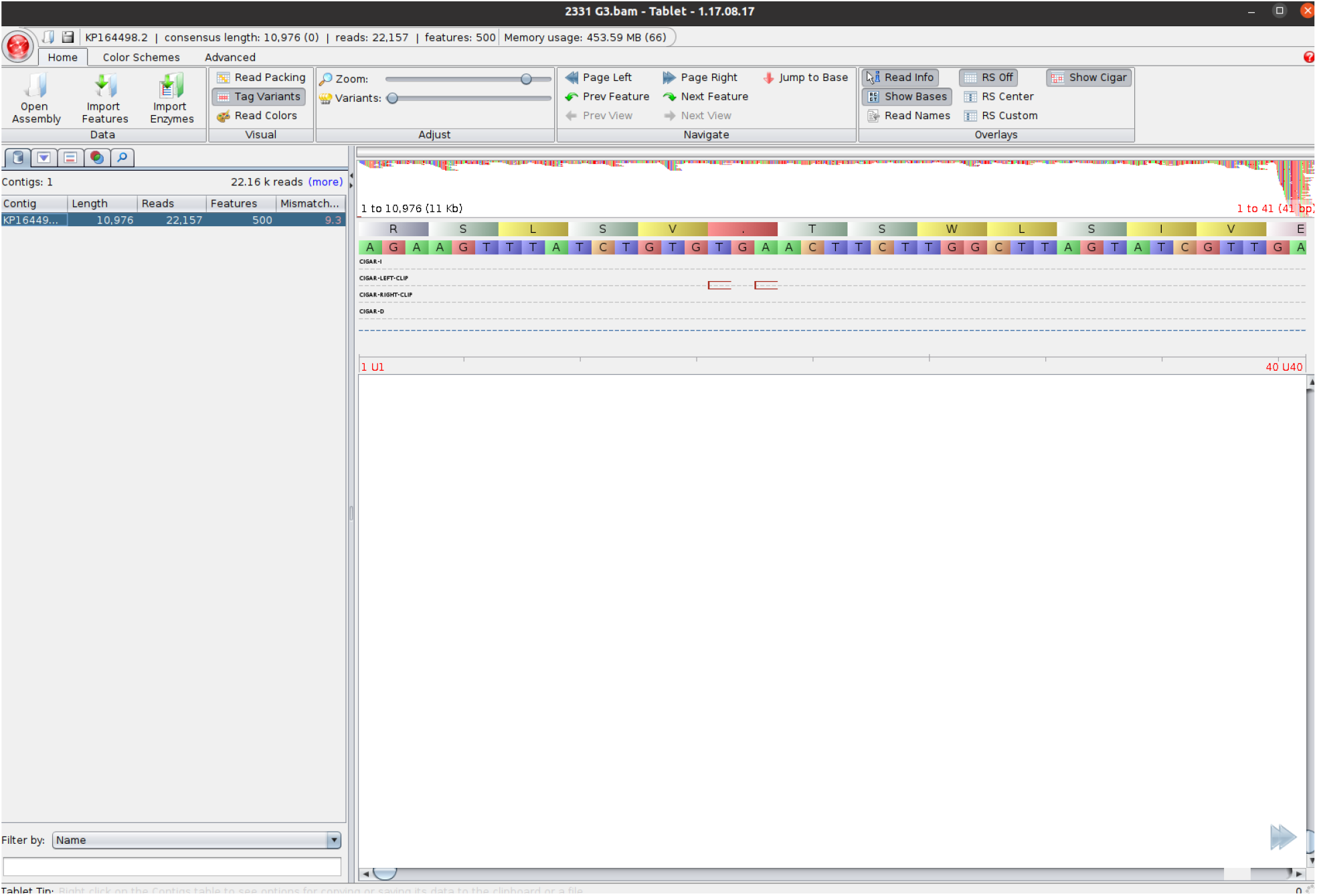
JEV positive sample 2331 processed with the Twist CVRP on MiSeq, contig aligned against the JEV G3 genome and viewed in Tablet

**Supplementary Figure 3.**
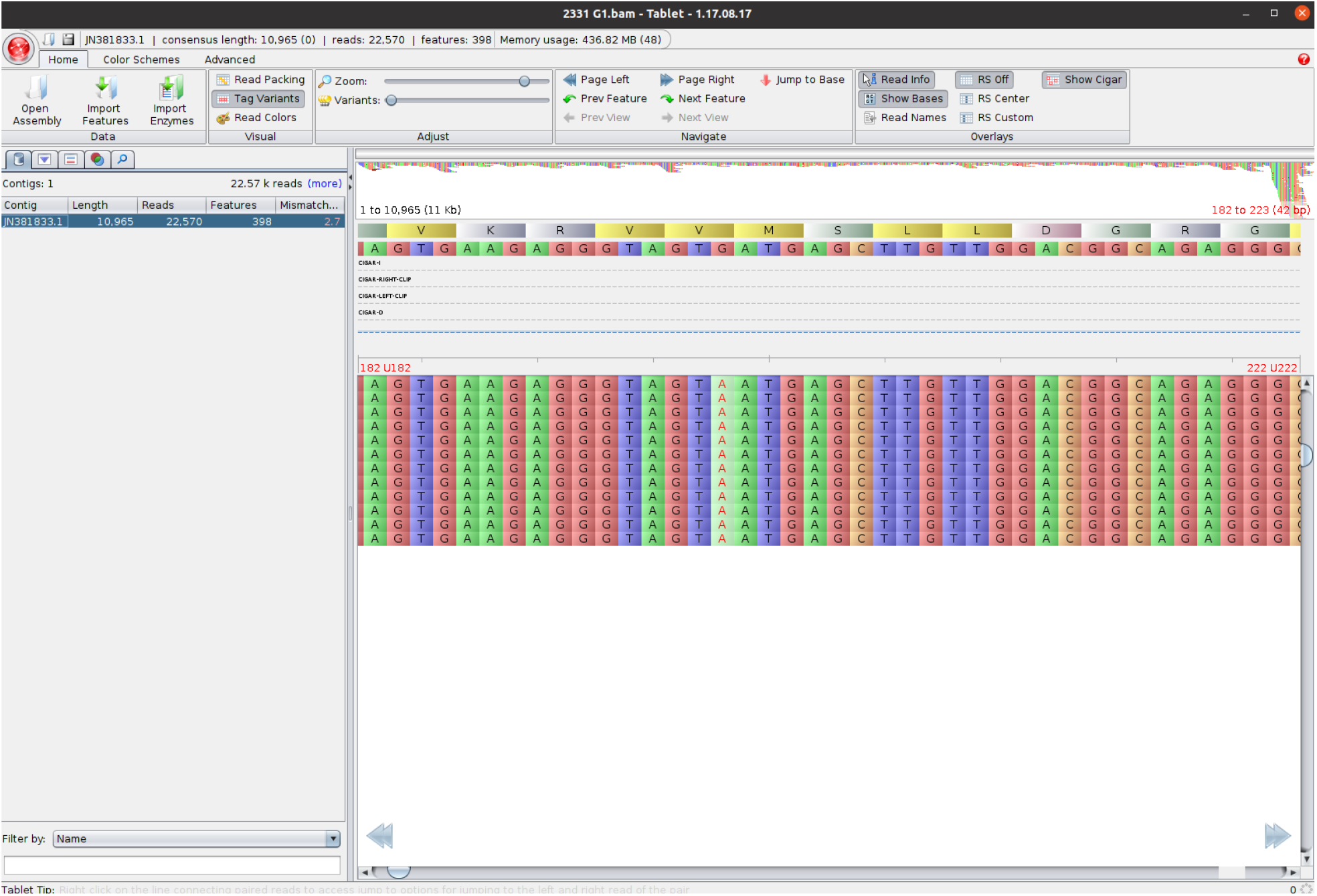
JEV positive sample 2331 processed with the Twist CVRP on MiSeq, contig aligned against the JEV G1 genome and viewed in Tablet

## CITATIONS

Agarwal, A., Parida, M., & Dash, P. K. (2017). Impact of transmission cycles and vector competence on global expansion and emergence of arboviruses. Reviews in Medical Virology, 27(5), e1941. https://doi.org/10.1002/rmv.1941

Amicizia, D., Zangrillo, F., Lai, P. L., Iovine, M., & Panatto, D. (2018). Overview of Japanese encephalitis disease and its prevention. Focus on IC51 vaccine (IXIARO®). Journal of preventive medicine and hygiene, 59(1), E99–E107. https://doi.org/10.15167/2421-4248/jpmh2018.59.1.962

Auerswald, H., Maquart, P.-O., Chevalier, V., & Boyer, S. (2021). Mosquito Vector Competence for Japanese Encephalitis Virus. Viruses, 13(6). https://doi.org/10.3390/v13061154

Calvert, A. E., Dixon, K. L., Delorey, M. J., Blair, C. D., & Roehrig, J. T. (2014). Development of a small animal peripheral challenge model of Japanese encephalitis virus using interferon deficient AG129 mice and the SA14-14-2 vaccine virus strain. Vaccine, 32(2), 258–264. https://doi.org/10.1016/j.vaccine.2013.11.016

Cao, L., Fu, S., Gao, X., Li, M., Cui, S., Li, X., Cao, Y., Lei, W., Lu, Z., He, Y., Wang, H., Yan, J., Gao, G. F., & Liang, G. (2016). Low Protective Efficacy of the Current Japanese Encephalitis Vaccine against the Emerging Genotype 5 Japanese Encephalitis Virus. PLOS Neglected Tropical Diseases, 10(5), e0004686. https://doi.org/10.1371/journal.pntd.0004686

Centers for Disease Control and Prevention. (2019). Japanese Encephalitis Vaccine | Japanese Encephalitis. https://www.cdc.gov/japaneseencephalitis/vaccine/index.html

Choi, M. B., Lee, W.-G., Kang, H. J., Yang, S.-C., Song, B. G., Shin, E.-H., & Kwon, O. (2017). Seasonal prevalence and species composition of mosquitoes and chigger mites collected from Daegu, Gunwi and Sangju in South Korea, 2014. Journal of Ecology and Environment, 41(1), 15. https://doi.org/10.1186/s41610-017-0030-7

Chrzastek, K., Lee, D., Smith, D., Sharma, P., Suarez, D. L., Pantin-Jackwood, M., & Kapczynski, D. R. (2017). Use of Sequence-Independent, Single-Primer-Amplification (SISPA) for rapid detection, identification, and characterization of avian RNA viruses. Virology, 509, 159–166. https://doi.org/10.1016/j.virol.2017.06.019

Eick-Cost, A. A., Hu, Z., Klein, T. A., Putnak, R. J., & Jarman, R. G. (2015). Seroconversion to Japanese Encephalitis Virus Among U.S. Infantry Forces in Korea. The American Society of Tropical Medicine and Hygiene, 93(5), 1052–1054. https://doi.org/10.4269/ajtmh.15-0307

Faizah, A. N., Kobayashi, D., Amoa-Bosompem, M., Higa, Y., Tsuda, Y., Itokawa, K., Miura, K., Hirayama, K., Sawabe, K., & Isawa, H. (2021). Evaluating the competence of the primary vector, Culex tritaeniorhynchus, and the invasive mosquito species, Aedes japonicus japonicus, in transmitting three Japanese encephalitis virus genotypes. PLOS Neglected Tropical Diseases, 14(12), 1–18. https://doi.org/10.1371/journal.pntd.0008986

Heng, L. (2018, June 18). Seqtk-1.3 (r106): Toolkit for processing sequences in FASTA/Q Formats. GitHub. https://github.com/lh3/seqtk

Hills, S. L., Lindsey, N. P., Fischer, M. (2019). Japanese Encephalitis - Chapter 4 - 2020 Yellow Book. Chapter 4 Travel - Related Infectious Diseases. https://www.nc.cdc.gov/travel/yellowbook/2020/travel-related-infectious-diseases/japanese-encephalitis

Hoke Jr., C. H. (2005). History of U.S. Military Contributions to the Study of Viral Encephalitis. Military Medicine, 170(suppl_4), 92–105. https://doi.org/10.7205/MILMED.170.4S.92

Infectious Disease Surveillance Center, National Institute of Infectious Disease (2017). Japanese encephalitis, Japan, 2007-2016. Infectious Agents Surveillance Report, 38(8). https://www.niid.go.jp/niid/images/idsc/iasr/38/450e.pdf

Kim H, Cha G-W, Jeong YE, Lee W-G, Chang KS, Roh JY, et al. (2015) Detection of Japanese Encephalitis Virus Genotype V in Culex orientalis and Culex pipiens (Diptera: Culicidae) in Korea. PLoS ONE 10(2): e0116547. doi:10.1371/journal.pone.0116547

Li, M.-H., Fu, S.-H., Chen, W.-X., Wang, H.-Y., Guo, Y.-H., Liu, Q.-Y., Li, Y.-X., Luo, H.-M., Da, W., Duo Ji, D. Z., Ye, X.-M., & Liang, G.-D. (2011). Genotype V Japanese Encephalitis Virus Is Emerging. PLOS Neglected Tropical Diseases, 5(7), e1231. https://doi.org/10.1371/journal.pntd.0001231

Lindahl, J., Chirico, J., Boqvist, S., Thu, H. T. V., & Magnusson, U. (2012). Occurrence of Japanese Encephalitis Virus Mosquito Vectors in Relation to Urban Pig Holdings. The American Society of Tropical Medicine and Hygiene, 87(6), 1076–1082. https://doi.org/10.4269/ajtmh.2012.12-0315

Lobigs, M., Pavy, M., Hall, R. A., Lobigs, P., Cooper, P., Komiya, T., Toriniwa, H., & Petrovsky, N. (2010). An inactivated Vero cell-grown Japanese encephalitis vaccine formulated with Advax, a novel inulin-based adjuvant, induces protective neutralizing antibody against homologous and heterologous flaviviruses. Journal of General Virology, 91(6), 1407–1417. https://doi.org/10.1099/vir.0.019190-0

Magoč, T., & Salzberg, S. L. (2011). FLASH: fast length adjustment of short reads to improve genome assemblies. Bioinformatics (Oxford, England), 27(21), 2957–2963. https://doi.org/10.1093/bioinformatics/btr507

Milne, I., Stephen, G., Bayer, M., Cock, P. J. A., Pritchard, L., Cardle, L., Shaw, P. D., & Marshall, D. (2013). Using Tablet for visual exploration of second-generation sequencing data. Briefings in Bioinformatics, 14(2), 193–202. https://doi.org/10.1093/bib/bbs012

Oliveira, A. R. S., Cohnstaedt, L. W., Noronha, L. E., Mitzel, D., McVey, D. S., & Cernicchiaro, N. (2020). Perspectives Regarding the Risk of Introduction of the Japanese Encephalitis Virus (JEV) in the United States. Frontiers in Veterinary Science, 7. https://doi.org/10.3389/fvets.2020.00048

Pan, X.L., Liu, H., Wang, H.Y., Fu, S.H., Liu, H.Z., Zhang, H.L., Li, M.H., Gao, X.Y., Wang, J.L., Sun, X.H., Lu, X.J., Zhai, Y.G., Meng, W.S., He, Y., Wang, H.Q., Han, N., Wei, B., Wu, Y.G., Feng, Y., Yang, D.J., Wang, L.H., Tang, Q., Xia, G., Kurane, I., Rayner, S., Liang, G.D. (2011). Emergence of Genotype I of Japanese Encephalitis Virus as the Dominant Genotype in Asia. Journal of Virology, 85(19). https://doi.org/10.1128/JVI.00825-11

Ratto-Kim, S., Yoon, I.-K., Paris, R. M., Excler, J.-L., Kim, J. H., & O’Connell, R. J. (2018). The US Military Commitment to Vaccine Development: A Century of Successes and Challenges. Frontiers in Immunology, 9. https://doi.org/10.3389/fimmu.2018.01397

Richards, E. E., Masuoka, P., Brett-Major, D., Smith, M., Klein, T. A., Kim, H. C., Anyamba, A., & Grieco, J. (2010). The relationship between mosquito abundance and rice field density in the Republic of Korea. International Journal of Health Geographics, 9(1), 32. https://doi.org/10.1186/1476-072X-9-32

Ricklin, M. E., Garcìa-Nicolàs, O., Brechbühl, D., Python, S., Zumkehr, B., Posthaus, H., Oevermann, A., & Summerfield, A. (2016). Japanese encephalitis virus tropism in experimentally infected pigs. Veterinary Research, 47(1), 34. https://doi.org/10.1186/s13567-016-0319-z

Sabin, A. B. (1947). EPIDEMIC ENCEPHALITIS IN MILITARY PERSONNEL: Isolation of Japanese B Virus on Okinawa in 1945, Serologic Diagnosis, Clinical Manifestations, Epidemiologic Aspects and Use of Mouse Brain Vaccine. Journal of the American Medical Association, 133(5), 281–293. https://doi.org/10.1001/jama.1947.02880050001001

Sanborn, M. A., Wuertz, K. M., Kim, H.-C., Yang, Y., Li, T., Pollett, S. D., Jarman, R. G., Berry, I. M., Klein, T. A., & Hang, J. (2021). Metagenomic analysis reveals Culex mosquito virome diversity and Japanese encephalitis genotype V in the Republic of Korea. Molecular Ecology, 30(21), 5470–5487. https://doi.org/10.1111/mec.16133

Shabardina, V., Kischka, T., Manske, F., Grundmann, N., Frith, M. C., Suzuki, Y., & Makalowski, W. (2019). NanoPipe—a web server for nanopore MinION sequencing data analysis. GigaScience, 8(2), giy169. https://doi.org/10.1093/gigascience/giy169

Shao, N., Li, F., Nie, K., Fu, S. H., Zhang, W. J., He, Y., Lei, W. W., Wang, Q. Y., Liang, G. D., Cao, Y. X., & Wang, H. Y. (2018). TaqMan Real-time RT-PCR Assay for Detecting and Differentiating Japanese Encephalitis Virus. Biomedical and Environmental Sciences, 31(3), 208–214. https://doi.org/10.3967/bes2018.026

Seo, H.J., Kim, H.C., Klein, T.A., Ramey, A.M., Lee, J.H., Kyung, S.G., Park, J.Y., Cho, Y.S., Cho, I.S., Yeh, J.Y. (2013). Molecular Detection and Genotyping of Japanese Encephalitis Virus in Mosquitoes during a 2010 Outbreak in the Republic of Korea. PLoS ONE. 8(2): e55165. https://doi.org/10.1371/journal.pone.0055165

Solomon, T., Ni, H., Beasley, D. W. C., Ekkelenkamp, M., Cardosa, M. J., & Barrett, A. D. T. (2003). Origin and evolution of Japanese encephalitis virus in southeast Asia. Journal of Virology, 77(5), 3091–3098. https://doi.org/10.1128/jvi.77.5.3091-3098.2003

World Health Organization. (2019). Japanese Encephalitis. Japanese Encephalitis. https://www.who.int/news-room/fact-sheets/detail/japanese-encephalitis

Wright, M. S., Stockwell, T. B., Beck, E., Busam, D. A., Bajaksouzian, S., Jacobs, M. R., Bonomo, R. A., & Adams, M. D. (2015). Sispa-Seq for rapid whole genome surveys of bacterial isolates. Infection, Genetics and Evolution, 32, 191–198. https://doi.org/10.1016/j.meegid.2015.03.018

Zhang, B., He, Y., Xu, Y., Mo, F., Mi, T., Shen, Q. S., Li, C., Li, Y., Liu, J., Wu, Y., Chen, G., Zhu, W., Qin, C., Hu, B., & Zhou, G. (2018). Differential antiviral immunity to Japanese encephalitis virus in developing cortical organoids. Cell Death & Disease, 9(7), 719. https://doi.org/10.1038/s41419-018-0763-y

